# Identification of novel regulators of dendrite arborization using cell type-specific RNA metabolic labeling

**DOI:** 10.1101/2020.09.28.316455

**Authors:** Mohamed Y. Aboukilila, Josephine D. Sami, Jingtian Wang, Whitney England, Robert C. Spitale, Michael D. Cleary

**Affiliations:** Department of Molecular and Cell Biology, Quantitative and Systems Biology Graduate Program, University of California, Merced, CA, USA; Department of Pharmaceutical Sciences and Department of Chemistry, University of California, Irvine, CA, USA

## Abstract

Obtaining neuron transcriptomes is challenging; their complex morphology and interconnected microenvironments make it difficult to isolate neurons without potentially altering gene expression. Multidendritic sensory neurons (md neurons) of *Drosophila* larvae are commonly used to study peripheral nervous system biology, particularly dendrite arborization. We sought to test if EC-tagging, a biosynthetic RNA tagging and purification method that avoids the caveats of physical isolation, would enable discovery of novel regulators of md neuron dendrite arborization. RNAs were biosynthetically tagged by expressing CD:UPRT (a nucleobase-converting fusion enzyme) in md neurons and feeding 5-ethynylcytosine (EC) to larvae. Tagged RNAs were subsequently purified and used for RNA-sequencing. Reference RNA was prepared in a similar manner using 5-ethynyluridine (EUd) to tag RNA in all cells and negative control RNA-seq was performed on “mock tagged” samples to identify non-specifically purified transcripts. Differential expression analysis identified md neuron enriched and depleted transcripts. Three candidate genes encoding RNA-binding proteins (RBPs) were tested for a role in md neuron dendrite arborization. Loss-of-function for the m6A-binding factor Ythdc1 did not cause any dendrite arborization defects while RNAi of the other two candidates, the poly(A) polymerase Hiiragi and the translation regulator Hephaestus, caused significant defects in dendrite arborization. This work provides an expanded view of transcription in md neurons and a technical framework for combining EC-tagging with RNA-seq to profile transcription in cells that may not be amenable to physical isolation.

## Introduction

Neuron development requires regulation of gene expression at the transcriptional and post-transcriptional levels. *Drosophila* peripheral nervous system (PNS) neurons provide a useful model for investigating these mechanisms. Sensory neurons of the larval PNS are classified according to dendrite morphology: external sensory and chordotonal neurons have a single dendrite, bipolar dendrite neurons have two unbranched dendrite projections, and multidendritic (md) neurons have more complex dendritic arborization. Md neurons innervate the larval body wall and function as touch receptors, proprioceptors, thermoreceptors or nociceptors. Md neurons have proven useful for investigating the molecular mechanisms that control dendrite arborization [1]. Foundational work used a mutagenesis screen to identify genes that regulate dendrite arborization [2]. Others have taken a reverse genetics approach and tested candidates through RNA-interference (RNAi) based on the functional properties of candidates [3, 4]. The ability to induce neuron-specific RNAi in *Drosophila* makes reverse genetics an attractive approach and the efficacy of candidate choice can be improved by selecting genes from neuron-specific transcriptome data.

Transcriptome profiling of dendritic arborization (da) neurons, a subclass of md neurons, has been performed using fluorescence-activated cell sorting (FACS) [5] or magnetic bead-based purification [6]. These studies used differential expression data to select candidates and demonstrated roles for those genes in da neuron dendrite arborization. One caveat of physical isolation is that neurons are removed from their natural environment and undergo processing prior to RNA extraction, possibly inducing transcriptional and post-transcriptional responses that do not reflect *in vivo* gene expression. An alternative approach is to use biosynthetic RNA tagging methods that do not require physical isolation and enrich for nascent and recently-transcribed mRNAs [7]. These methods use metabolic labeling, under *in vivo* conditions, to generate tagged RNAs in the cells of interest. Tagged RNAs are subsequently purified from total RNA of animals or tissues. We recently described a cell type-specific biosynthetic RNA tagging method called EC-tagging [8]. EC-tagging works via targeted expression of a nucleobase-converting fusion enzyme composed of cytosine deaminase and uracil phosphoribosyltransferase (CD:UPRT). Metazoans lack cytosine deaminase activity and have varying endogenous uracil phosphoribosyltransferase activity, depending on organism or cell type [9, 10]. Only cells expressing CD:UPRT are competent to convert the bioorthogonal base 5-ethynylcytosine (EC) into 5-ethynyluracil (via CD activity) and subsequently into 5-ethynyluridine monophosphate (via transgenic UPRT and endogenous pathways) which is ultimately incorporated into nascent RNAs. The 5-ethynyl group allows click-chemistry-based biotinylation of tagged RNAs and subsequent purification on streptavidin beads. We previously demonstrated the utility of EC-tagging in the *Drosophila* central nervous system (CNS) [8]. This initial work used a microarray platform to analyze transcriptomes of relatively large populations of neurons (the entire larval CNS and the mushroom body neurons). To further test the specificity and sensitivity of this technique and to discover novel regulators of dendrite arborization, here we combine EC-tagging with RNA-sequencing to generate md neuron transcriptome profiles.

## Results

### EC-tagging enriches for larval md neuron-specific transcripts

To identify genes transcribed in all md neurons, we used *Gal4^109(2)80^* [2] to drive *UAS-CD:UPRT* and fed 5EC to larvae from 72 - 84 hours after hatching. Based on fluorescent imaging of tagged RNA (data not shown), we estimate it takes a minimum of six hours before ingested 5EC is disseminated to and metabolized by target cells. The twelve-hour 5EC feeding is therefore expected to generate a population of tagged RNAs synthesized over approximately six hours. At the end of EC feeding, larval carcass (containing primarily muscle, epidermis and the peripheral neurons of interest) was dissected to remove CNS neurons that express *Gal4^109(2)80^*. A reference sample was prepared by feeding 5-ethynyluridine (5EUd) to stage-matched *UAS-CD:UPRT* larvae, in the absence of any Gal4. 5EUd is incorporated into RNA independent of CD:UPRT and thus provides a reference containing mRNAs transcribed in all cells over the same labeling period. As a negative control, we prepared “mock-tagged” samples in which larvae were not fed 5EC or 5EUd but were subjected to the same carcass dissection and RNA processing. The mock sample serves as a control for the stringency of the purification and allows identification of transcripts that may be purified independent of EC-tagging. This type of mock reference has proven useful in other biosynthetic RNA labeling experiments [11, 12].

For all three sample types (5EC-tagged, 5EUd-tagged and mock-tagged), equal amounts of total RNA were biotinylated and applied to streptavidin beads. RNA captured on the beads was directly used to prepare sequencing libraries. The number of mapped reads per sample agreed with the expected yield of tagged RNA: 5EUd-tagging (expected high yield, RNA tagging in all cells) gave 48 – 56.6 million reads, 5EC-tagging (expected low yield, RNA tagging only in rare md neurons) gave 2.7 – 7.7 million reads, and mock-tagging (background) gave 0.11 – 0.13 million reads. 5EC-tagged biological replicates and 5EUd-tagged biological replicates had a high degree of RNA-seq correlation, while the correlation for mock-tagged replicates was much lower (**Fig. S1**). To identify transcripts enriched in md neurons, we performed DE-seq analysis comparing 5EC-tagged RNA (EC-RNA) and 5EUd-tagged RNA (ref-RNA). This differential expression analysis was performed using two versions of the ref-RNA: 1) the complete RNA-seq dataset and 2) a randomly generated subset of 2.7 million reads. The random subset matches the read depth of the lowest yield EC-RNA library, thus controlling for possible sample size effects.

DE-Seq identified 937 enriched transcripts and 236 depleted transcripts (minimum two-fold difference and adjusted p-value < 0.05). To address potential non-specific purification among the enriched transcripts, we compared transcript levels between EC-RNA and mock-RNA. This comparison identified 85 transcripts with no significant enrichment compared to mock-RNA, thus reducing the list of md neuron transcripts to 852. Similar background correction for depleted transcripts is not possible since transcripts that are rare or absent in md neurons are expected to be low in EC-RNA and mock-RNA. In the background-corrected dataset, known md neuron transcripts are among the most significantly enriched and known muscle-specific transcripts are among the most significantly depleted **(Fig. 1)**. The top 60 enriched or depleted genes are listed in **Figure 2** and the complete DE-seq results are provided in **Table S1**. EC-RNA compared to the subset ref-RNA yielded 571 enriched transcripts and 130 depleted transcripts (minimum two-fold difference and adjusted p-value < 0.05) and the background correction procedure removed 39 transcripts from the enriched list (**Table S1)**. Enriched and depleted transcripts were similar regardless of the type of ref-RNA used (commonly enriched transcripts are denoted in **Table S1**). We also performed gene ontology (GO) analysis on the complete ref-RNA DE-Seq results (**Fig. 3**) and the subset ref-RNA DE-Seq results (**Fig. S2**). Both approaches yielded multiple neuron-specific GO categories including “peripheral nervous system development” and “dendrite morphogenesis”. The “synaptic growth at neuromuscular junction” GO category reflects the fact that enriched genes in this category function in the synapses of motor neurons and md sensory neurons. Unexpected GO categories, “dorsal closure” and “border follicle cell migration”, reflect the fact that many neurite growth or morphogenesis genes (e.g. *stathmin, shot, kay, shn, aop*) are also involved in these processes. In addition to the EC-tagged data, we performed GO analysis on DE-Seq results comparing mock-RNA to the complete or subset ref-RNA. Importantly, the mock-tagged DE-Seq data did not result in any significant GO category enrichment.

**Figure 1.**
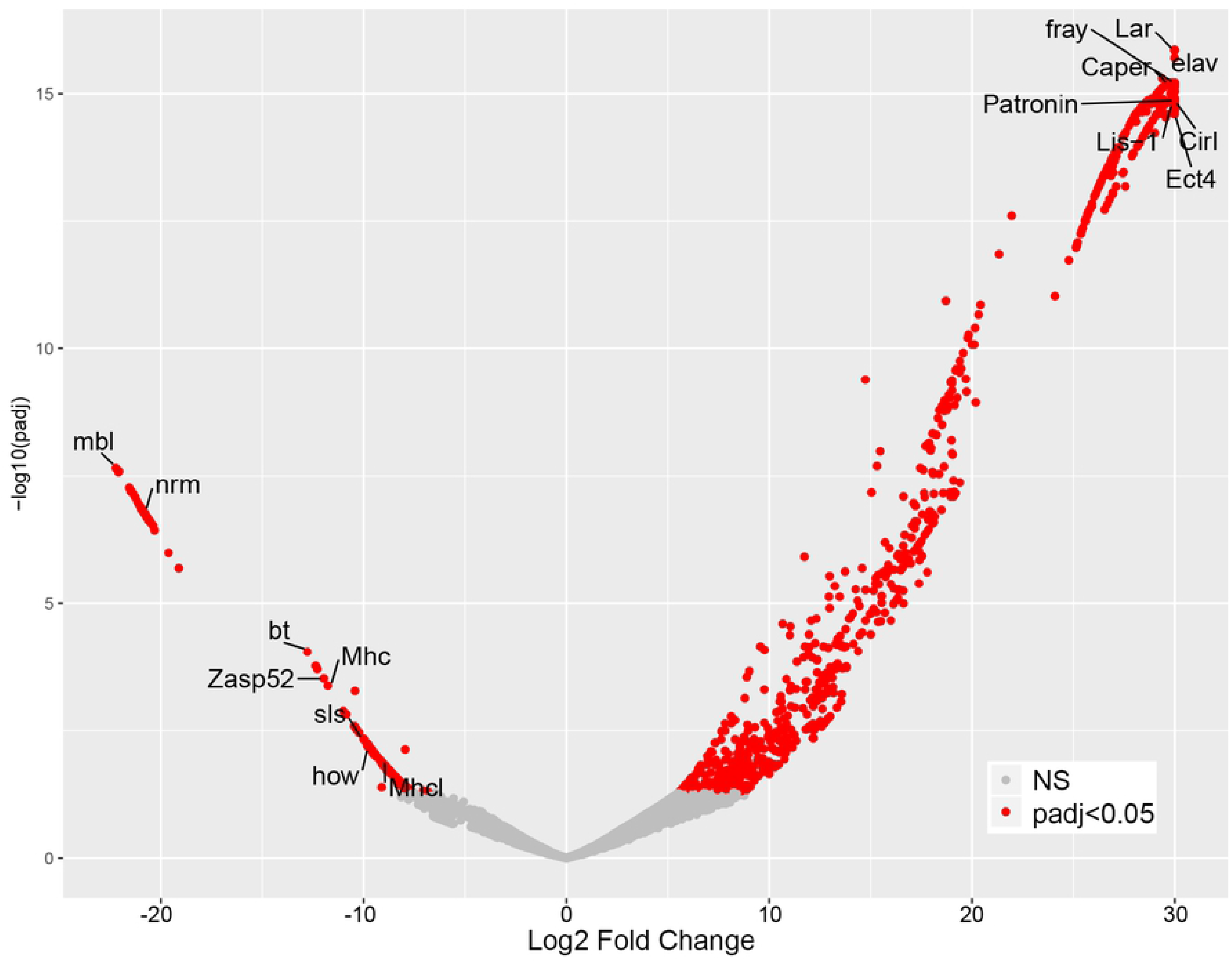
Identification of md neuron enriched and depleted transcripts. Volcano plot of DE-Seq results showing the log_2_ fold change for EC-tagged TPM / EUd-tagged TPM (x-axis) and the adjusted p-value for each transcript (y-axis). NS = not significant. Select genes with known md neuron expression (enriched side of plot) and known muscle expression (depleted side of plot) are labeled.

**Figure 2.**
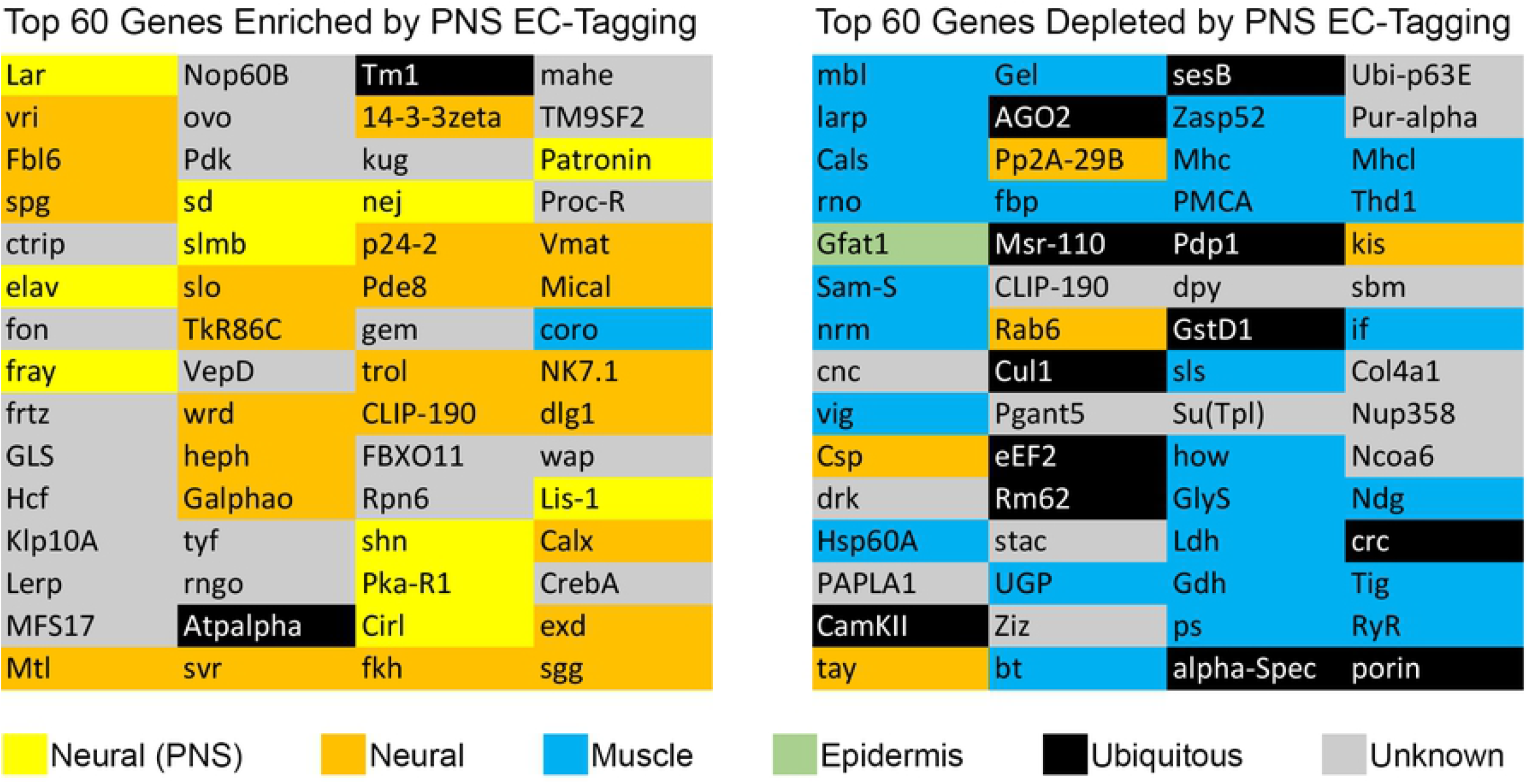
Top md neuron enriched and depleted transcripts. Only named genes are included, genes known only by “CG number” were excluded. The primary tissue expression pattern is color coded. Tissue expression was determined by searching Flybase annotations and published literature. “Neural (PNS)” indicates known expression in larval md neurons. “Neural” indicates expression in the central nervous system at any stage. “Muscle” indicates expression in any type of muscle at any stage. “Epidermis” indicates expression in epidermis at any stage. “Ubiquitous” indicates either widespread expression or evidence of expression in both neurons and muscle. “Unknown” indicates the literature do not support assignment to any of the other categories.

**Figure 3.**
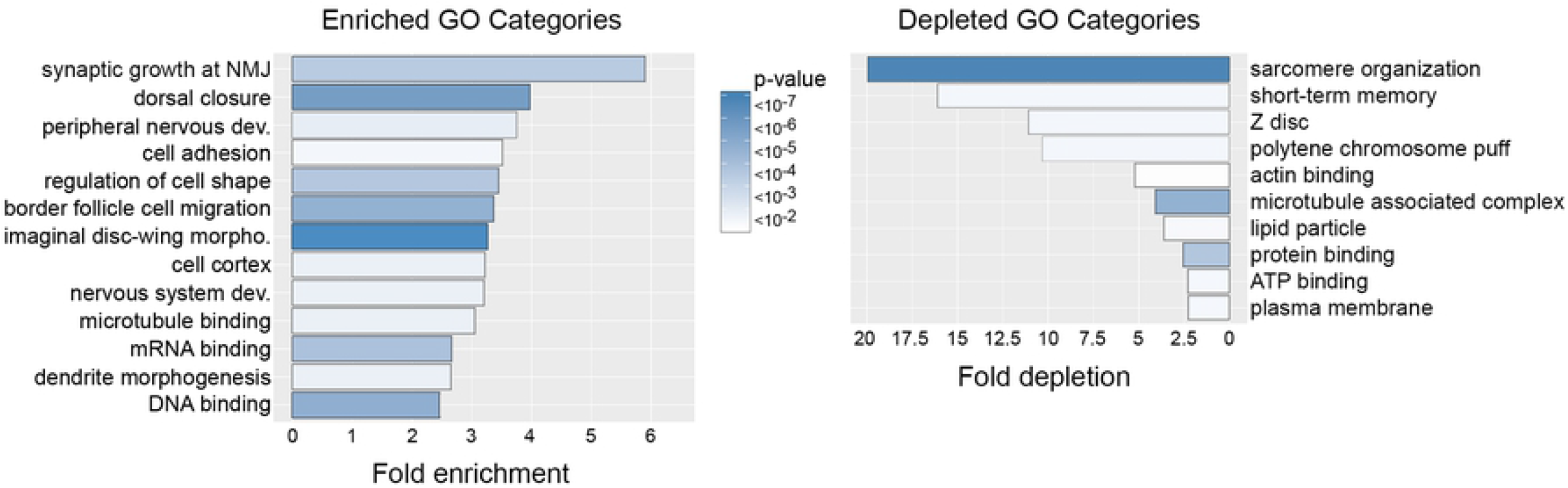
Gene ontology of enriched and depleted transcripts. GO categories over-represented among EC-RNA enriched and depleted genes. Observed / expected value = frequency of category genes in EC-RNA / frequency in the *Drosophila* genome. Heatmap = Bonferroni-corrected P-values. NMJ = neuromuscular junction.

Given the agreement between EC-RNA compared to the full or subset ref-RNA, we focused subsequent analyses on the background-corrected EC-RNA versus full ref-RNA. In addition to the genes listed in Figure 2, this dataset contains many previously described md neuron genes (according to Flybase annotations and associated references), including transcription regulators (*ab, ttk, stan, cnc, jim, kay, gro*), RNA-binding factors (*bel, Caper, Fmrp, sqd, stau, rump*), ion channels (*Piezo, SK*), signal receptors and transducers (*EcR, Egfr, Rac1, spin, puc*), and cytoskeletal factors (*spas, shot).* We also identified multiple transcripts encoding general regulators of neurotransmission, including *Frq1, brp, Csp, and Rim.* While differences in experimental design and target cell populations limit the validity of broad comparisons between our data and prior transcriptome studies, we selected two gene lists from prior studies to compare with our enriched gene set. Iyer et al. identified 40 transcription factor genes enriched in class I and/or class IV da neurons compared to whole larvae [5]. Our enriched gene set contains 6 of these 40 genes, a moderate but significant over-representation (Fisher’s exact test comparing representation in the EC-RNA dataset to representation in the *Drosophila* genome, p-value = 0.03). Hattori et al. identified 24 genes expressed in class I and/or class IV da neurons for which RNAi caused dendritic arborization phenotypes [6]. Our md neuron enriched gene set contains 10 of these 24 genes, a significant over-representation (Fisher’s exact test, p-value = 5 x 10^-6^).

### Candidate testing identifies novel regulators of md neuron dendrite arborization

We next sought to test the function of novel candidate genes in md neuron dendrite arborization. We focused on genes encoding mRNA-binding proteins (a significantly enriched GO category, Fig. 3)) based on our interest in post-transcriptional control of mRNA processing [13]. Out of 20 enriched RNA-binding proteins (RBPs), 10 (*rump, fus, bru1, sqd, stau, shep, elav, Fmr1, BicD*, and *Caper*) have previously described functions in md neuron dendrite arborization (based on Flybase annotations or a RNAi screen of RBPs [4]). The RNAi screen of Olesnicki et al. [4] tested 7 additional RBPs that we identified as md neuron-enriched but did not detect any dendrite arborization defects, suggesting these RBPs have functions irrelevant to dendrite morphogenesis. We focused on the remaining 3 RBPs with no available md neuron information: *ythdc1* (also known as *YT521-B), hiiragi (hrg*), and *hephaestus* (*heph*). Ythdc1 is a nuclear-localized m6A binding protein that regulates alternative splicing [14]. Hiiragi is a poly(A) polymerase that acts on nascent mRNAs and regulates poly(A) tail length of cytoplasmic mRNAs in oocytes [15]. Hephaestus is a polypyrimidine tract binding protein that represses translation of *oskar* in oocytes [16]. We selected a known md neuron RBP enriched in our dataset, *Caper*, as a positive control. Caper regulates alternative splicing and is required for proper dendrite arborization of the ddaC class IV da neuron [17].

We crossed *UAS-RNAi* lines to *Gal4^477^, UAS-mCD8::GFP* [4]. The membrane-tethered GFP allowed us to measure the number of dendrite branch terminals and dendritic field size in ddaC neurons of late L3 larvae. *Gal4^477^* expression begins in newly differentiated class IV da neurons of embryos and continues throughout larval development [18]. *Gal4^477^, UAS-mCD8::GFP* crossed to wildtype served as a negative control. *Caper* RNAi caused a significant increase in dendrite branch terminals but no increase in dendritic field size since the terminal branches tended to be short and tightly clustered (**Fig. 4**). These *Caper* RNAi results are very similar to the previously described *Caper* loss-of-function phenotype [17]. For the analysis of *ythdc1*, we used a previously described loss of function mutant [19] combined with *Gal4^477^, UAS-mCD8::GFP* as well as a *ythdc1* RNAi line. Neither *Ythdc1* loss of function (**Fig. 4**) nor ythdc1 RNAi (data not shown) affected the number of dendrite branch terminals or field coverage. In contrast, *Heph* RNAi caused a significant decrease in the number of branch terminals and field size (**Fig. 4**). *Hrg* RNAi also caused a significant decrease in branch terminals and field size but the termini / field size value was not significantly different from controls (**Fig. 4**). These results suggest that *heph* knockdown primarily affects branching, resulting in fewer terminals and reduced dendritic field size, while *hrg* knockdown primarily affects dendrite growth without directly affecting branching.

**Figure 4.**
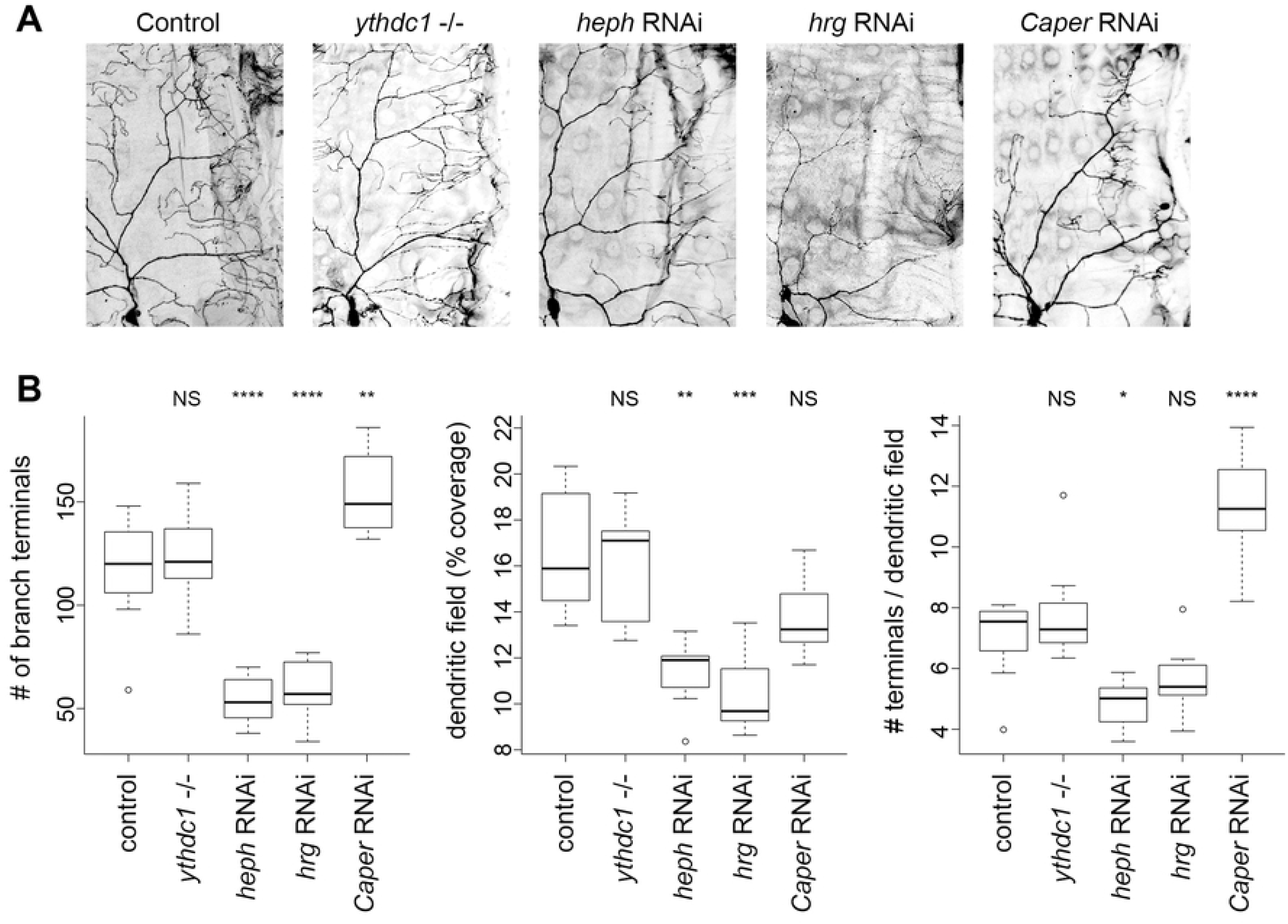
Identification of novel regulators of md neuron dendrite arborization. **A.** Dendrite arborization in control and mutant class IV ddaC neurons. A representative image is shown for each genotype. The same region was analyzed in all cases, with the cell body positioned at lower left. Larva anterior is to the left and dorsal is at the top. **B.** Quantification of dendrite arborization. Within the window applied to all neurons (as shown in panel A), the number of branch terminals were counted and the total dendrite length was traced and quantified (pixels) then divided by the total pixels of the window to calculate the dendritic field (% coverage). The number of terminals per neuron was divided by the dendritic field % coverage to calculate # terminals / dendritic field. Data were analyzed using ANOVA with Tukey HSD post-test. P-value code: **** ≤ 0.0001, *** ≤ 0.001, ** ≤ 0.01, * ≤ 0.05.

## Discussion

We selected larval multidendritic sensory neurons for transcription profiling for two reasons: 1) this cell population presents a good target for testing the specificity and sensitivity of EC-tagging and 2) md neurons have previously been used to identify post-transcriptional regulators of dendrite arborization and we reasoned that EC-tagging would enable identification of additional genes in this category. Our prior EC-tagging work [8] used a microarray platform and focused on relatively large populations of cells (all neurons of the central nervous system, mushroom body neurons); in contrast, this work demonstrates that EC-tagging can be combined with RNA-seq and is sensitive enough to identify transcripts in a rare neuronal population. Through the use of EUd-tagged and mock-tagged references, we obtained significant enrichment of expected md sensory neuron transcripts and a list of potential novel regulators of md neuron development and function.

Neuronal metabolic RNA labeling experiments have revealed that profiling newly transcribed mRNAs can reveal gene expression dynamics that are missed by more traditional steady-state measurements [20, 21]. Analysis of mRNAs synthesized over a relatively short period likely aided our detection of transcripts that have a high rate of decay, such as transcription factors, signaling factors, and RNA binding proteins (classes of genes we have shown to encode low stability mRNAs [13]). For example, in the embryonic nervous system, mRNAs encoding the three RBPs we selected for analysis (*ythdc1, heph*, and *hrg*) all have below average half-lives [13]. As previously described [7], metabolic RNA tagging approaches are most effective when comparing purified target cell RNA to a reference generated by metabolic tagging in all cell types of the starting material (tissue or whole animal). For this reason, any genes transcribed in more abundant cells of the carcass are unlikely to be identified as md neuron-enriched in this study. For example, Iyer et al. identified the transcription factors *Poxm* and *Hand* as transcribed in da neurons but these genes are also transcribed in larval muscle and are abundant in our EUd ref-RNA. EC-tagging may be best suited for the discovery of differentially transcribed genes while physical isolation methods such as FACS are better suited for defining complete transcriptomes. Depending on experiment goals, the potential for a less comprehensive transcriptome profile generated by EC-tagging should be weighed against the risks of perturbing gene expression through the use of physical isolation.

RNA-binding proteins have previously been shown to play important roles in md neuron dendrite arborization [4]. Given the significance of this class of post-transcriptional regulators in dendrite arborization, we prioritized the analysis of RBPs from among all the candidate regulators of md neuron development identified in this study. Olesnicky et al. used a comprehensive RNAi-based screen to identify a large number of RBPs that affect class IV da neuron dendrite morphogenesis. We identified three additional md neuron-enriched RBPs of interest: *ythdc1, hrg* and *heph.* Knockdown of *hrg* and *heph* resulted in dendrite arborization defects while *ythdc1* loss of function and knockdown had no effect. The lack of a dendrite arborization phenotype in the *ythdc1* mutants may indicate this is a false positive (not expressed in md neurons). It is difficult to test this possibility without a Ythdc1 antibody, but the widespread expression of Ythdc1 in CNS neurons [14] suggests a general neural function. An alternative explanation is that Ythdc1 controls splicing of transcripts that are irrelevant to dendrite arborization. *Drosophila* motor neurons that lack m6A, the RNA modification recognized by Ythdc1, do not have growth or patterning defects but do have a moderate increase in the number of synaptic boutons and active zones per bouton [14]. Loss of Ythdc1 in md neurons may similarly affect synapses or synaptic activity, phenotypes that would not be detected in our analysis. The *hrg* RNAi phenotype may be explained by Hrg’s interaction with Orb, an ortholog of human cytoplasmic polyadenylation element binding protein 1. Hypomorphic alleles of *orb* cause dendrite arborization defects in class IV da neurons [22], with decreased branching and decreased field size similar to what we observed in *hrg* RNAi neurons. Hrg and Orb may regulate cytoplasmic polyadenylation in md neurons, likely targeting mRNAs encoding regulators of dendrite growth. Heph may affect dendrite arborization via its repression of *oskar (osk*) translation. Oskar is necessary for proper localization of *nanos* mRNA in class IV da neurons and *osk* loss of function decreases dendrite branching [23], similar to the phenotype we observe in *heph* RNAi neurons. *Osk* mRNA is transported along dendrites [23] and Heph likely represses *osk* translation during transport, as it does in oocytes [16]. In this model, *heph* knockdown may cause a phenotype similar to *osk* loss of function due to altered Osk distribution in dendrites. Confirming these predicted interactions and further defining mechanisms by which Hrg and Heph control dendrite arborization will be important areas of future investigation.

## Materials and Methods

### Drosophila genetics

The following lines were obtained from the Bloomington *Drosophila* Stock Center: Oregon-R-P2 (wildtype) (stock # 2376), *Gal4^477^, UAS-mCD8::GFP* (stock # 8746), *UAS-ythdc1{RNAi}* (stock #34627), UAS-hrg{RNAi} (stock # 33378), UAS-heph{RNAi} (stock # 55655). For EC-tagging, *Gal4^109(2)80^* (stock # 8769) was combined with *UAS-CD:UPRT* on the 3^rd^ chromosome (stock # 77120) to make the stable line *Gal4^109(2)80^; UAS-CD:UPRT.* The ythdc1 loss of function mutant, *YT521-B[NP2]/ TM6C −1*, was provided by Dr. Eric Lai.

### EC-tagging, RNA purification and sequencing library preparation

5-ethynylcytosine was synthesized as previously described [8]. Biological replicates were prepared by carrying out 5EC, 5EUd, or mock feeding, carcass dissections and RNA processing independently. Larvae were reared at 25°C and fed 1 mM 5EC or 5EUd from 72 – 84 hours after hatching. Total carcass RNA was extracted using Trizol. For each treatment, duplicate 20 μg RNA samples were biotinylated using Click-iT Nascent RNA Capture reagents (ThermoFisher) and purified on Dynabeads MyOne Streptavidin T1 magnetic beads (ThermoFisher) as previously described [8]. After the final wash, beads were combined with NuGen Ovation Universal RNA-Seq reagents, following the manufacturer’s protocol beginning at first strand cDNA synthesis. Primer annealing and cDNA synthesis was performed in a heated lid thermomixer to ensure beads did not settle. After cDNA synthesis, beads were washed three times with 500 μl 1X PBS, discarding the supernatant each time. Beads were resuspended in 50 μl RNaseA/T1/H elution mix (1X RNAse H buffer, 12.5 mM D-biotin, RNase A/T1 cocktail (0.1 U/μl), RNase H (0.1 U/μl)) and incubated at 37°C for 30 minutes at 1,000 rpm in a thermomixer. The reaction was stopped by adding 1 μl of DMSO and heating at 95°C for 4 minutes. Beads were collected on a magnet and the supernatant was mixed with DNA binding buffer and applied to a Zymogen DNA Clean and Concentrator-5 column to purify first-strand cDNA. Following purification, the duplicate samples were combined into a single tube and the volume was reduced to 10 μl using a SpeedVac concentrator. The samples were then used to make second-strand cDNA according to the NuGen Ovation Universal RNA-Seq protocol, including adapter ligation and ribosomal RNA depletion using a *Drosophila-specific* AnyDeplete rRNA primer mixture. Libraries were amplified and purified according to the NuGen protocol and quality was assessed using an Agilent Bioanalyzer DNA high-sensitivity chip.

### RNA-sequencing and bioinformatics

Sequencing was performed on a HiSeq 2500. Sequence data were trimmed using *Trimmomatic* prior to mapping to the *Drosophila melanogaster* cDNA transcriptome (BDGP6) using *kallisto.* Differential expression analysis was performed using *DESeq2.* The EUd-RNA reference subset was obtained using the *shuf* command in UNIX to randomly select 2.7 million reads from the EUd-RNA BAM files.

### Imaging and quantification of dendrite morphology

Dendrite morphology was analyzed in wandering larval stages. Larval fillet preparations were fixed using formaldehyde and stained with rat anti-mCD8 (ThermoFisher) at 1:100 followed by Alexa Fluor 488 anti-rat secondary antibody (ThermoFisher) at 1:200. Imaging was performed using a Zeiss LSM 880 confocal microscope. The number of branch terminals and total dendrite length were quantified in Z-series projections. Branch terminal counting and dendrite tracing was performed using Adobe Photoshop and dendrite length (pixels) were quantified using Zeiss Zen Blue software. Statistical significance was determined using ANOVA with Tukey HSD post-test.

## Acknowledgements

We thank Dr. Eric Lai for stocks used in this study. Stocks obtained from the Bloomington Drosophila Stock Center (NIH P40OD018537) were also used in this study. We thank members of the UC Irvine Genomic High Throughput Facility for technical assistance. Imaging data were collected with a confocal microscope acquired through the National Science Foundation MRI Award Number DMR-1625733. The authors acknowledge funding from the National Institutes of Health (5R21MH116415 to M.D.C. and R.C.S.).

## Supporting Information

Fig. S1. RNA-seq correlation between biological replicates.

Fig. S2. GO category enrichment from EC-RNA versus subset ref-RNA DE-Seq.

Table S1. DE-Seq data for EC-RNA versus full and subset ref-RNA, DE-Seq data for EC-RNA versus mock-RNA.

